# Mutations, Recombination and Insertion in the Evolution of 2019-nCoV

**DOI:** 10.1101/2020.02.29.971101

**Authors:** Aiping Wu, Peihua Niu, Lulan Wang, Hangyu Zhou, Xiang Zhao, Wenling Wang, Jingfeng Wang, Chengyang Ji, Xiao Ding, Xianyue Wang, Roujian Lu, Sarah Gold, Saba Aliyari, Shilei Zhang, Ellee Vikram, Angela Zou, Emily Lenh, Janet Chen, Fei Ye, Na Han, Yousong Peng, Haitao Guo, Guizhen Wu, Taijiao Jiang, Wenjie Tan, Genhong Cheng

## Abstract

**Background:** The 2019 novel coronavirus (2019-nCoV or SARS-CoV-2) has spread more rapidly than any other betacoronavirus including SARS-CoV and MERS-CoV. However, the mechanisms responsible for infection and molecular evolution of this virus remained unclear.

**Methods:** We collected and analyzed 120 genomic sequences of 2019-nCoV including 11 novel genomes from patients in China. Through comprehensive analysis of the available genome sequences of 2019-nCoV strains, we have tracked multiple inheritable SNPs and determined the evolution of 2019-nCoV relative to other coronaviruses.

**Results:** Systematic analysis of 120 genomic sequences of 2019-nCoV revealed co-circulation of two genetic subgroups with distinct SNPs markers, which can be used to trace the 2019-nCoV spreading pathways to different regions and countries. Although 2019-nCoV, human and bat SARS-CoV share high homologous in overall genome structures, they evolved into two distinct groups with different receptor entry specificities through potential recombination in the receptor binding regions. In addition, 2019-nCoV has a unique four amino acid insertion between S1 and S2 domains of the spike protein, which created a potential furin or TMPRSS2 cleavage site.

**Conclusions:** Our studies provided comprehensive insights into the evolution and spread of the 2019-nCoV. Our results provided evidence suggesting that 2019-nCoV may increase its infectivity through the receptor binding domain recombination and a cleavage site insertion.

**One Sentence Summary:** Novel 2019-nCoV sequences revealed the evolution and specificity of betacoronavirus with possible mechanisms of enhanced infectivity.

## Introduction

A new coronavirus, named the Novel Coronavirus 2019 (2019-nCoV or SARS-CoV-2), has emerged and been transmitted to the human population^1^. The outbreak originating from Wuhan, China began in December of 2019 and currently has, as of Feb 23^th^, 2020, over 77,000 confirmed cases globally and over 2400 fatalities^2^. The current infection has spread outwards from China to 29 other countries or regions including South Korea, Japan, Thailand, Singapore, Vietnam, Taiwan, Nepal, and the United States^3^. Series of pneumonia cases from Wuhan linked to this virus, with symptoms such as fever, dry cough, and dyspnea^4,5^. Most patients had no significant upper respiratory tract symptoms, suggesting that target cells are located lower in the airway. Consequently, the virus was isolated and sequenced from epithelial cells of the lower respiratory tract^5^.

Coronaviruses are enveloped, positive sense, single-stranded RNA viruses^6^ that undergo rapid mutation and recombination^7,8^, of its receptor binding domain (RBD) to adapt to a large pool of species^9–14^. Analysis of the 2019-nCoV shows that it is a novel betacoronavirus belonging to the lineage B (subgenus: *sarbecovirus*) which also includes Severe Acute Respiratory Syndrome Coronavirus (SARS-CoV)^4^. The current 2019-nCoV genome is most phylogenetically similar to Bat/SARS/RaTG13/Yunnan strain, which was first isolated in 2013 in Yunnan, China^15^. Recent studies hinted that pangolin-CoV may be a possible intermediate hosts candidate for the 2019-nCoV^16,17^. Recently, this novel CoV has been confirmed to use the same cell entry receptor, ACE2, as SARS-CoV^16^. Coronaviruses (CoV) are capable of transmitting among different host species via shifting tropism and variable receptor targeting^9–13^. These characteristics are mediated by changes in the receptor binding domain (RBD) of the spike surface glycoprotein^12,14^. Similar to other well-known enveloped viruses, CoV initiates infection through fusion of this spike protein with the host cell membrane^18^. Their spike protein is comprised of S1 and S2 subunits, which are responsible for host receptor recognition and membrane fusion, respectively^12,13,18,19^. 2019-nCoV has recently been confirmed to use the human ACE2 as receptor^20^.

Although the fatality rate of 2019-nCoV is lower and the symptoms are milder than those of SARS, the transmissibility of 2019-nCoV appears to be higher^7^. The molecular mechanisms responsible for such rapid transmission and spread of the 2019-nCoV still remain elusive. Here, we have analyzed 120 2019-nCoV genome sequences including 11 novel strains isolated from patients to determine the viral evolution and divergence. Our studies suggest that 2019-nCoV may increase its infectivity through the receptor binding domain recombination and a furin or TMPRSS2 cleavage site insertion.

## Results

The first cases of human-to-human 2019-nCoV infection occurred mid-December 2019, To date, the total confirmed cases and confirmed death surpassed SARS by a magnitude within a shorter period of time^1^. In an attempt to associate the pathologic property of 2019-nCoV with specific virulence factor, we compared the epidemiological information with sequencing data obtained from the WHO^2^. We compared the rate of infection for 2019-nCoV to that of the most recent betacoronavirus outbreaks, SARS in November 2002 and MERS in September 2012 (Fig 1A.). 2019-nCoV appears to be transmitted much more quickly than SARS and MERS. To date, there are more confirmed 2019-nCoV cases than that of the whole 2002 SARS outbreak. Coronavirus sequences have been published frequently and consistently over the last 18 years, which included SARS strains isolated from different countries during 2002 SARS outbreak, sporadic CoV strains mainly reported in China, MERS strains isolated from middle east countries such as Saudi Arabia and United Arab Emirates (Fig 1B). To further understand the evolution of the betacoronaviruses and track the mutations accumulated with 2019-nCoV, we have collected and sequenced 11 full-length 2019-nCoV genome sequences from new patients identified in multiple Chinese cities including Wuhan (Fig 1C). The phylogenetic analysis indicated that 11 new strains of 2019-nCoV clustered together with other 2019-nCoV strains and were more homologous to the bat CoV RaTG13 strain than human SARS, MERS and other CoVs (Fig S1A). At the amino acid levels, they only had a few random substitutions at positions with consensus sequences identical to the corresponding amino acid sequences in human and bat SARS (Fig S1B).

**Figure 1.**
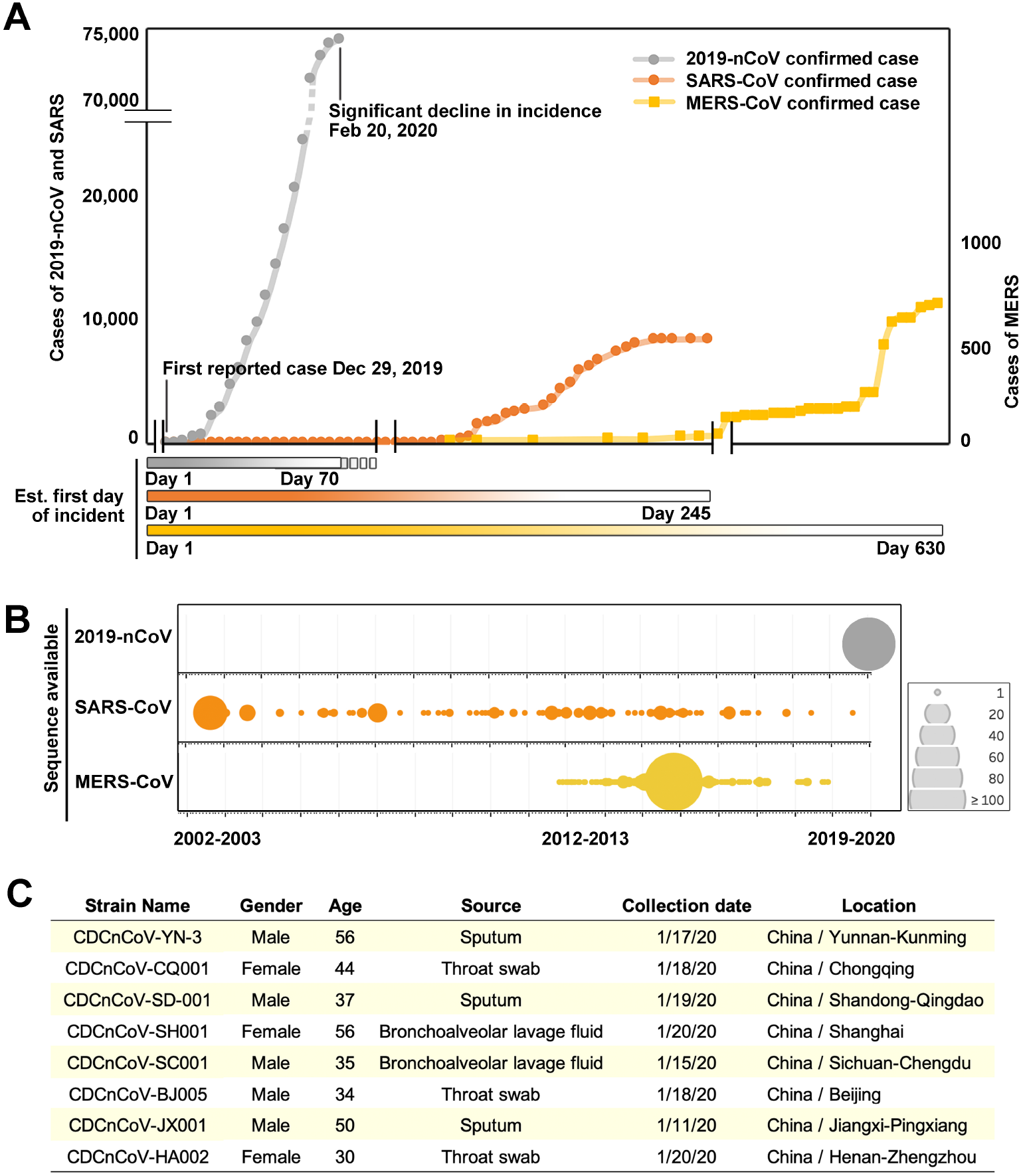
**(A)** Data of total confirmed cases during SARS (orange line), MERS (yellow line) and 2019-nCoV (grey line) epidemic. **(B)** Number of sequences published and available in public domains in Asia and Middle East from Dec 2002 to Feb 2020. All panels are current as of Feb 24^th^, 2020. **(C)** Information about samples taken from patients infected with 2019-nCoV. nCoV is the 2019 novel coronavirus. SARS-CoV is severe acute respiratory syndrome coronavirus. MERS-CoV is Middle East respiratory syndrome coronavirus.

To identify novel inherited mutations, we used SARS-CoV-2 strain (EPI_ISL_402125) as root to construct the phylogenetic tree for all 120 available complete genomes of the novel coronavirus from GISAID (updated February 18^th^, 2020). Considering potential sequencing errors at both ends, the genome sequence variations among different strains of 2019-nCoV isolated from patients located in different cities were low considering only several mutations in about 30kb genome per isolate. Based on the nucleotide positions 8517 and 27641, the 2019-nCoV strains can be divided into two major groups (Figs 2A and S2). All the group 1 strains have thymine at 8517 and cytosine at 27641, which are same as corresponding nucleotides in SARS, whereas the group 2 stains have cytosine at 8517 and thymine at 27641 (Figs 2A and S3). Epidemiological data G1 and G2 strains revealed that the collection date and location of the earliest G1 strain (EPI_ISL_406801) was January 5^th^, 2020 in Wuhan, whereas the earliest G2 strain was isolated in December 24^th^, 2019 in Wuhan (Fig 2A). The existence of both genetic groups in the same city indicated co-circulation, but evolved convergently at the early outbreak. Within each group, we also observed additional shared mutations added to multiple strains. Based on these potentially inheritable mutations and the identifying times and locations, we generated a mutation tree map to track individual shared mutations and show the relationships among different isolates (Fig 2A and S2). For example, five strains identified in Guangdong from Jan. 10-15 all share the same mutation in nucleotide position 28578 on the background of group 1 might be transmitted by the same person. The similar strain may transmit to three patients identified in Japan on Jan. 29-31 with additional mutation at nucleotide position 2397 and to a patient identified in USA on Jan. 22 with additional mutation at nucleotide position 10818. The G10818T is very interesting as it is shared by several independent strains in both group 1 and group 2, which will lead to L3606F amino acid substitution within the orf1ab polyprotein (Fig S3). It is not clear if the common mutation both in group 1 and group 2 strains at 10818 site has any growth advantage but pangolin and bat CoV have L3606V substitution at the same position (Fig S3). When the group 1 and group 2 strains were placed onto geographic map (Fig 2B), it seems that both groups have been transmitted to most countries and regions with reported 2019-nCoV cases with few exceptions, suggesting these two groups are rapidly transmissible.

**Figure 2.**
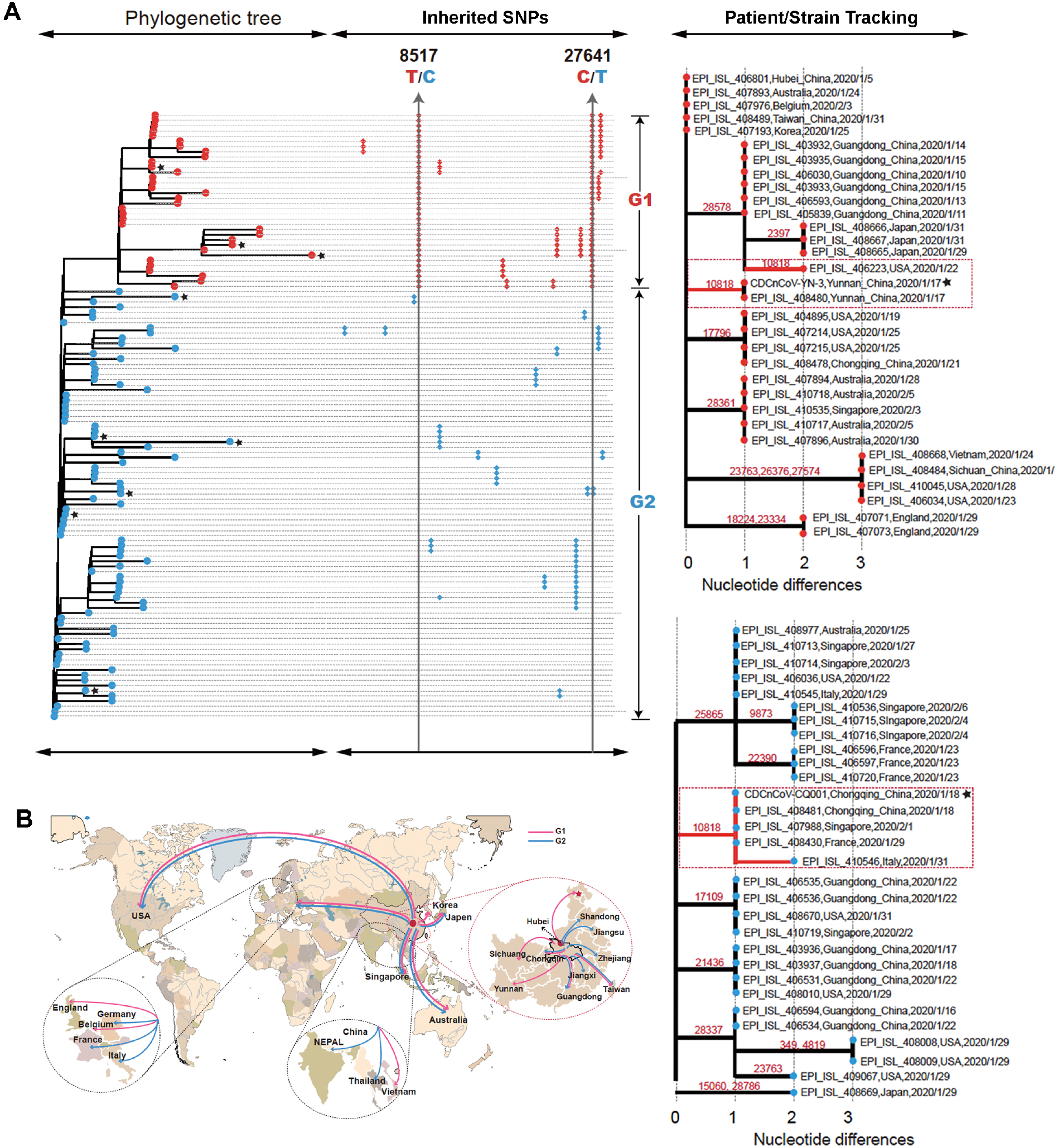
**(A)** Sequence alignment of 120 full-length genomes of 2019-nCoV including 11 newly reported genomes (highlighted by stars), ~30,000 base pairs in length, nucleotide substitutions to an early sequenced strain EPI_ISL_402125 as the root of phylogenetic tree. Two sub-groups were coloured in blue and red. The trailing dots on the right represent the SNPs in the viral genome. The first group (G1) possesses 8517T and 27641C. The second group (G2) possesses 8517C and 27641T. All genomes were clustered using ML method. Inherited SNPs that were share between multiple strains were highlighted. The divergent evolution of G1 and G2 was identified by tracking individual shared mutations. The horizontal axis represents the difference between reference sequence and the node strain. **(B)** Geographical of the spread of different genetic groups of 2019-nCoV. The red and blue lines represent predicted transmission pathways with G1 and G2 strains, respectively.

The most closely related strain, betacoronavirus RaTG13, was isolated from *Rhinolophus affinis*^15^ (Fig 3A). We performed additional phylogenetic analyses using nucleotide sequences for specific viral proteins such asorf1a, spike, matrix and nucleocapsid (Fig. 3B), and found the same close relationship of the RaTG13 strain and other bat SARS-like CoV strains. We further estimated that the divergences of most proteins between 2019-nCoV and RaTG13 happened between 2005 and 2012 whereas those between human SARS and bat SARS-like CoV happened between 1990-2002 (Fig 3B). When the full-length spike protein sequences were compared, 2019-nCoV shares 39% sequence homology to human and bat SARS as compared to MERS or other CoV at 29%. Interestingly, we found that 2019-nCoV and pangolin-CoV share near identical amino acid sequence in the RBD (aa 315-550 region) of the spike protein, but not for RaTG13 (Figs 3A and 3D). To confirm this finding, we have compared the pangolin-CoV reported by Liu *et al* with previously isolated but unpublished pangolin-CoV sequences (Figure S4). Based on the alignment and phylogenetic analysis, we found that the consensus sequence of 2019-nCoV has the highest identity to BetaCoV/pangolin/Guangdong/P2S/2019 (EPI_ISL_410544), whereas additional mutations and indels were found in the pangolin-CoV strains isolated in the Guangxi province. Both BetaCoV/pangolin/Guangdong/P2S/2019 and Pangolin-CoV were isolated from Guangdong province of China in 2019. Next we examined the phylogenetic relationship of the consensus spike protein sequence of 2019-nCoV against 25 representative CoV strains including Hu-CoV, SARS and MERS and five new pangolin-CoV strains using a ML method (Figure S4). The results demonstrate that the Spike protein of 2019-nCoV is likely derived from Pangolin-CoV but not RaTG13, all of which could be in the same lineage as BAT_SARS-CoV/bat-SL-CoVZC542. The fact that while 2019-nCoV is most homologous to RaTG13 in overall genome structure the RBD of the spike protein is most homologous to Pangolin-CoV, suggests the possible recombination between RaTG13 like and Pangolin-CoV like strains happens in the evolution of 2019-nCoV. We have manually examined all amino acid mutations in the genome. Due to the presence of the unsequenced regions in the Pangolin-CoV sequence, we did not include any positions flanking these regions. We found that when comparing Pangolin-CoV and 2019-nCoV, in addition to the RBD, regions in nsp14 and 15 also shares consecutive sequences (Fig 3D).

**Figure 3.**
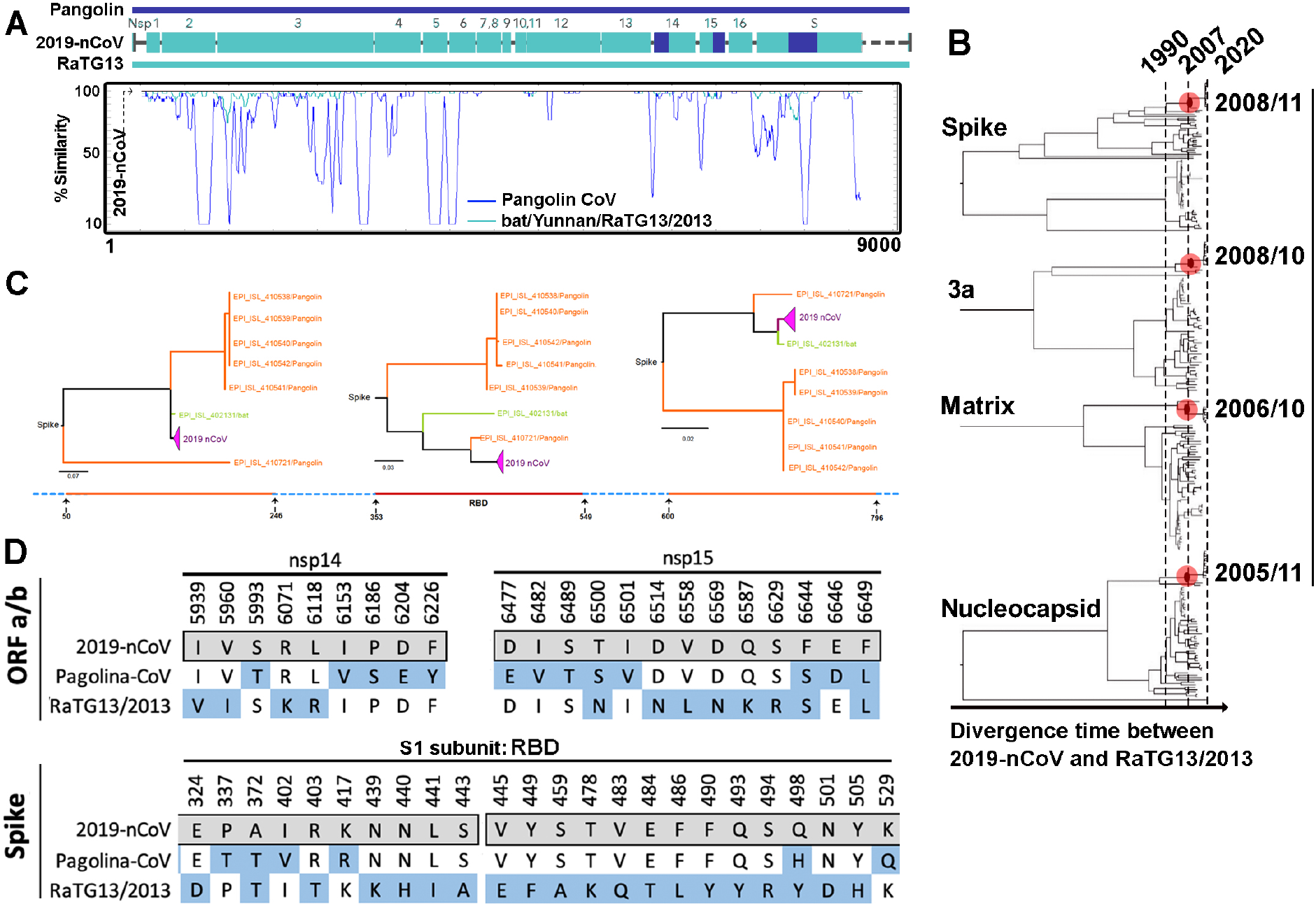
**(A)** Amino acid homology graph of Pangolin-CoV (blue line) and RaTG13/2013 (green line) against 2019-nCoV. Conserved receptor binding domain (RBD) highlighted in yellow. **(B)** Schematic phylogenetic trees based on the molecular clock analysis for the 2019-nCoV with the orf1a, spike, matrix and nucleocapsid genes. The phylogenetic clades that contain the 2019-nCoV, the SARS-like bat CoVs and the SARS viruses in human and the other hosts are colored by red, violet and light blue lines, respectively. The SARS-like bat CoV Yunnan-RaTG13 strain was highlight by the green line. The Divergence time in the molecular clock analysis between 2019-nCoV and RaTG13 were labelled. **(C)** Phylogenetic analysis of the aligned spike protein sequences from represented betacoronavirus strains using the ML method based on the JTT matrix-based model on MEGA v7.0. Alignment was performed using MUSCLE. 2019-nCoV, RaTG13 and Pangolin-CoV were highlighted with arrow. **(D)** Amino acid substitutions of Pangolin-CoV, 2019-nCoV and RaTG13. ORFa/b and spike proteins encoded by Pangolin-CoV and Bat/Yunnan/RaTG13 were aligned against 2019-nCoVs using MUSCLE. Sites flanking an unidentified amino acid were excluded.

To further evaluate the 2019-nCoV relationship with other SARS CoVs, we analyzed the RBM of the 2019-nCoV and different human/bat SARS viruses and observed that they can be clearly divided into two distinct clades (Fig 4A). The clade I viruses include 2019-nCoV, pangolin CoVs, 12 of bat SARS (bat SARS CoV I) such as RaTG13, and human SARS (Table S1). The clade II viruses include 49 of bat SARS viruses (bat SARS CoV II) such as ZXC21 and ZC45, which share about 90% overall nucleotide and amino acid identity as the 2019-nCoV (Table S2). The major difference between these two clades is that the RBM of the clade II viruses has regions with 5 and 13-14 amino acids shorter than that of the clade I viruses (Fig. 4B). Previous structural analysis have demonstrated that the 13-14 amino acid region of the SARS RBM forms a distinct loop structure, which is stabilized with a disulfide bond between two cysteine residues^9,15,16,21^. Although the amino acid sequences of 2019-nCoV within this loop region are very different from those of human SARS, the two cysteine residues are conserved (Fig. 4B). Interestingly, all the CoV viruses known to use ACE2 as entry receptor are within the clade I, including the 2019-nCoV, which can also infect cells through ACE2 receptor-based on recent studies^22^ (Fig. S5). On the other hand, there is no report on using ACE2 as entry receptor for the clade II viruses, despite their overall genome sequence homology with 2019-nCoV. Therefore, our studies not only emphasize the critical role of the RBM in determining the specificity of entry receptors but also raise the question on how homologous strains of betacoronavirus switch tropism through mutations such as indel or recombination in the RBM.

**Figure 4.**
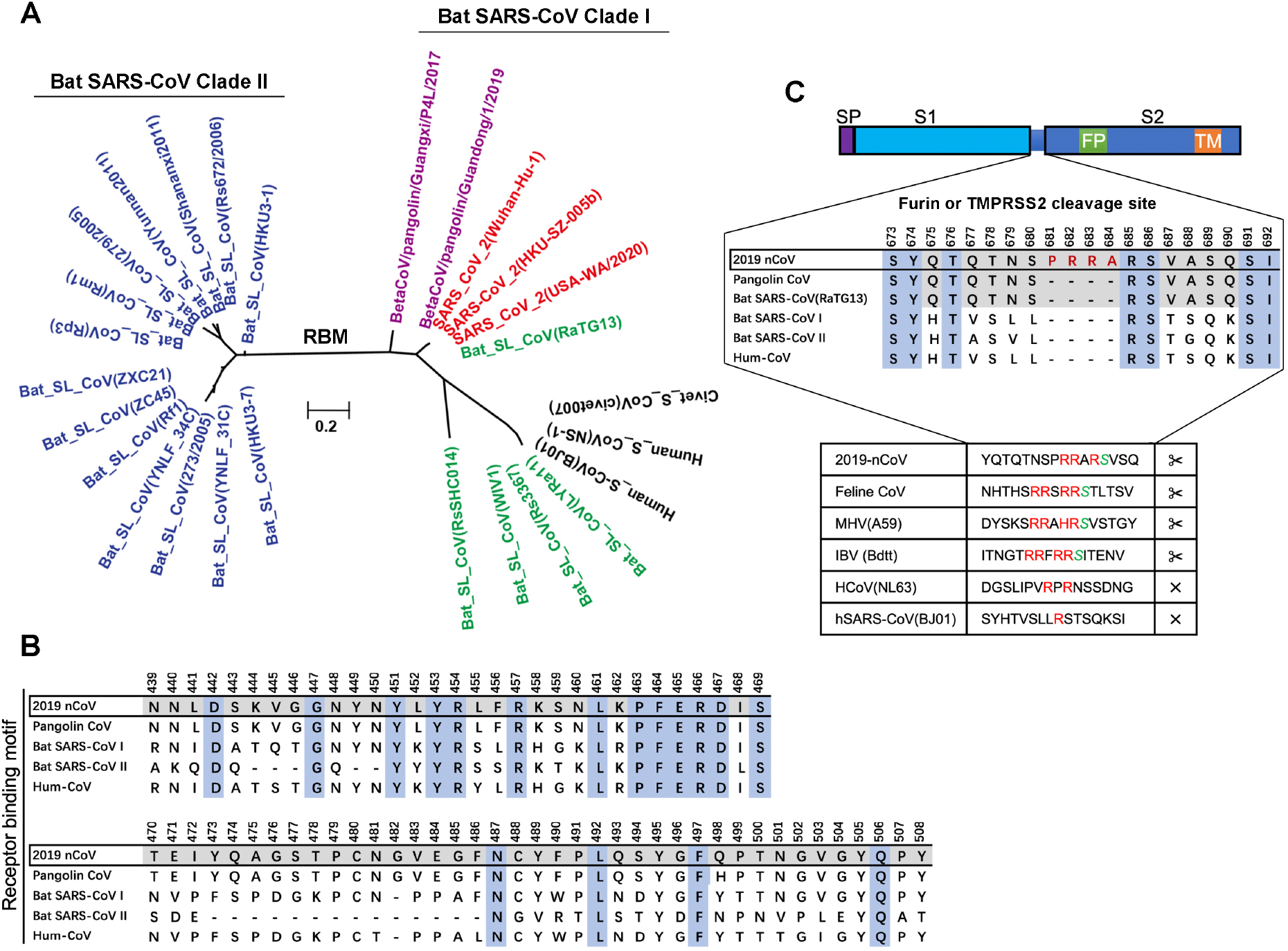
**(A)** Molecular Phylogenetic analysis of receptor binding motifs of 2019-nCoV, human/civet SARS virus and bat SARS-like virus. The evolutionary history was inferred by using the Maximum Likelihood method based on the JTT matrix-based model conducting in MEGA7 software. The scale bar represents the number of substitutions per site. **(B)** Specific amino acid variations in the RBM of the spike proteins of the 2019-nCoV and Bat SARS-CoV sub-lineages. **(C)** Coronavirus Spike proteins and their potential furin or TMPRSS2 cleavage sites between S1 and S2 domain. Insertion mutation at amino acid position 681 of 2019-nCoV spike protein and its sequence comparison with human and bat SARS-like CoVs. The multiple alignment was conducted by using the Clustal in MEGA7 software with default parameters. Furin consensus target sites were labeled with red; residues immediately downstream of the cleavage site are italicized and highlighted in green.

Our study also revealed that the 2019-nCoV has a unique four amino acid insertion (681-PRRA-684) within the spike protein, or within nucleotide position 23,619-23,632 (Fig 4C). Interestingly, such an insertion (PRRA) within the spike protein of 2019-nCoV creates a potential cleavage site RRAR for the mammalian furin protein, which the consensus sequence is RXXR. The potential furin or TMPRSS2 cleavage site is inserted at the boundary between the S1 and S2 domains of the spike protein, and the first proline residue of the PRRA insertion may introduce a beta turn into the polypeptide chain. To understand the uniqueness of this insertion, we performed sequence comparison using represented SARS-CoV strains from human, civet and bat. Our study showed that this insertion is unique to the 2019-nCoV (Figure 4C and S6). When compared to other CoV family members, we found that similar insertion has been identified and are located in the structural boundary between the S1 and S2 domains of the spike protein (Figure 4C).

## Discussion

The 2019-nCoV is still spreading rapidly from Wuhan to different cities in China and other countries, at a magnitude faster than SARS and MERS^4^. We have systematically analyzed and tracked the genome mutations among 120 different strains of 2019-nCoV. Although most substitutions are *de novo* mutations, we have identified multiple SNPs shared by different strains of 2019-nCoV. Due to the low mutation rate and large genome capital, we hypothesis that these shared mutations are inherited SNPs from large group of populations, likely transmitted from people to people. Based on this assumption, we have generated the SNP tree to track mutation and spread of 2019-nCoV. Interestingly, the 2019-nCoV stains we have analyzed fall into two distinct groups with different nucleotide polymorphisms at positions 8782 and 28144. Although it remains to be determined if these two distinct groups of 2019-nCoV were evolved before or after transmission from animal to human, both groups were first detected in wuhan and then spread to different regions in China and multiple countries. By combining additional inheritable mutations in subsequent generations of 2019-nCoV with the times and locations of the patient sample collections, we can trace possible viral transmission pathways.

2019-nCoV as a member of the betacoronaviruses, shares similar genome structures as bat SARS, human SARS, MERS with nucleotide identity over 88%, 79%, about and 50%,respectively^23^. Its closest relative is the Beta CoV RaTG13, isolated from bat in Yunnan province, China, in 2013, which shares more than 96% identical nucleotides throughout the genome of over 30 kb^15,24^. Our evolution clock analysis estimated that 2019-nCoV diverged from RaTG13 and human SARS-CoV at about 12 and 30 years ago, respectively. Beside point mutations, there is also a potential evidence of recombination as a mechanism for the evolution of 2019-nCoV. We found that 2019-nCoV shares high identity with RaTG13 throughout the genome except for the RBD of spike protein, which is closer to Pangolin-CoV isolated in Guangdong^17^. Based on these findings we hypothesized that recombination in the RBD of the spike protein may happen between RaTG13 and Pangolin-CoV like strains during the evolution of 2019-nCoV.

Bat is believed to be the original host for the 2019-nCoV but its intermediate host before transmitting to human is not known whereas civets and camels are widely considered as the intermediate hosts for human SARS and MERS, respectively^22,25–27^. Human SARS viruses are known to use the ACE2 while MERS use DPP4 as their receptors to infect host cells^16,28^. Recent studies have indicated ACE2 as the entry receptor for 2019-nCoV^29^ although other host cell factors such as TMPRSS2 are likely involved^30^. When we used the receptor binding motifs to conduct phylogenic analysis, we could separate 2019-nCoV, human and bat SARS into two distinct clades. Interestingly, all the viruses known to use ACE2 as the entry receptor are within the clade I whereas all the bat SARS viruses which do not use ACE2 entry receptors belong to clade II. Therefore, we predict that the clade I CoV viruses including the 2019-nCoV can while clade II CoV cannot infect host cells through ACE2. Based on the available genome sequences, there are many more clade II bat CoV (over 49 stains) than clade I bat CoV (about 12 strains). Some of the clade II bat CoV such as ZXC21 and ZC45 are more homologous to the 2019-nCoV than the clade I bat CoV and human SARS. It would be interesting to find out if homologous betacoronavirus can switch tropism through recombination in the RBM.

We have also identified a unique four amino acid (PRRA) insertion between S1 and S2 domains of the spike protein, which may function as a furin or TMPRSS2 cleavage site. It has been shown that CoV may undergo a protease cleavage to trigger the virus-cell membrane fusion^10,14^. This flexibility in priming and triggering the fusion machinery greatly modulates the viral pathogenicity and tropism of different coronaviruses^18,19^. However, such protease cleavage has not been detected in SARS-CoV^19^. Introducing a cleavage site into SARS-CoV resulted in spike protein cleavage and potentiated the membrane fusion activity^11^. In addition, introducing a cleaved spike protein into a SARS-CoV pseudotype virus allowed it to directly enter host cells^31^. Based on previous sequencing and structural analysis, the 2019-nCoV spike protein were predicted to interact with the ACE2 receptor to trigger the fusion with the host cell membrane and initiate infection^29^. Therefore, mutations or indels altering the S1-S2 subunits should significantly impact viral infection. We hypothesize that the PRRA insertion may render the spike protein to cleavage process, which triggers the viral fusion event.

Although the exact mechanisms responsible for such a high infection rates remains to be further investigated, our data on both recombination in the RBD and the unique furin or TMPRSS2 cleavage site insertion between the S1 and S2 domains of the spike protein may explain why the transmission of this newly emerged virus is significantly increased compared to the related beta-coronaviruses, including SARS and MERS. Further tracking the genome mutations with additional strains of 2019-nCoV isolated from patients in different locations at different time points will provide insights to understand the molecular evolution of this rapid spreading viruses. More importantly, comparison of protein sequence and structural differences between 2019-nCoV and other beta coronaviruses will provide insights to rapidly develop novel strategies to treat or prevent diseases associated with the novel emerging infectious viruses.

## Acknowledgment

This work was supported by NIH R01AI069120 and R01AI140718 grants, the National Key Plan for Scientific Research and Development of China (2016YFD0500301, 2016YFC1200200), CAMS Initiative for Innovative Medicine (CAMS-I2M, 2016-I2M-1-005), the National Natural Science Foundation of China (U1603126), the Central Public-Interest Scientific Institution Basal Research Fund (2016ZX310195, 2017PT31026 and 2018PT31016), China postdoctoral science foundation grant (2019M660548). We thank Dr. Fang Li for helpful discussion.

## Supplementary Materials for

### Experimental Procedures

#### Patients and samples

The whole-genome sequences of 2019-nCoV from 11 samples were generated by a combination of Sanger, Illumina, and Oxford nanopore sequencing. First, viral RNAs were extracted directly from clinical samples with the QIAamp Viral RNA Mini Kit, and then used to synthesize cDNA with the SuperScript III Reverse Transcriptase (ThermoFisher, Waltham, MA, USA) and N6 random primers, followed by second-strand synthesis with DNA Polymerase I, Large (Klenow) Fragment (ThermoFisher). Viral cDNA libraries were prepared with use of the Nextera XT Library Prep Kit (Illumina, San Diego, CA, USA), then purified with Agencourt AMPure XP beads (Beckman Coulter, Brea, CA, USA), followed by quantification with an Invitrogen Qubit 2.0 Fluorometer. The resulting DNA libraries were sequenced on either the MiSeq or iSeq platforms (Illumina) using a 300-cycle reagent kit. About 1 ·2–5 GB of data were obtained for each sample.

#### Genome and amino acid comparisons

The start and stop codons of each predicted gene were manually checked to ensure the completeness. We tried to infer the possible evolution pathway and track the cotransmission of 2019-nCoV. To avoid random mutations that may mislead the inference, we removed the substitution that appeared only once among all the sample genomes. Nucleotide alignment was performed using MUSCLE (version 2.2.25+), and base-pair comparison was performed using Basebybase (v1.0). Nucleotide substitutions were determine using a consensus sequence of 2019-nCoV. The comparison of amino acids within the 2019-nCoV were carried out. Additionally, for each ORF, the amino acid sequences of 2019-nCoV, Human SARS-CoV and Bat SARS-CoV were compared using the MUSCLE.

#### Phylogenetic analysis

Given the whole protein sequences corresponding for the 2019-nCoV, 2019-nCoVs were obtained from ViPR – Coronavirus and GISAID database on 18^th^ Feb, 2020. We performed a representative search with the related sequences of the betacoronaviridae virus using the BLAST (version 2.2.25+). Based on the blast results, we excluded the strains without the sample date, location, host and species information. Then the screened sequences were aligned by the MUSCLE. The sequences were removed fot that with no less than 28000bp or more than 100 unsolved nucleotides as ‘N’. At last, 109 public genomes of 2019-nCoV were kept. The phylogenetic tree for the whole genome was constructed using the Molecular Evolutionary Genetics Analysis (MEGA) (version 7.0)^32^ by the maximum likelihood method under the General Time Reversible (GTR) nucleotide substitution model, while the phylogenetic trees for the individual genes were constructed by the FigTree software v1.4.3.

For the spike protein, given the whole genomes of 2019 nCoVs (118 strains) and other SARS-like viruses isolated from Bat(1 strains) and Pangolin (6 strains), the genes was predicted by GeneMarkS (version 3.36). The predicted genes were then against the proteome of SARS-CoV by BLAST (version 2.2.25+). After that, Spike surface glycoprotein (S) were picked, and RBD domain in Spike was also discriminated. For the protein sequence of Spike genes, we got, multiple sequence alignment was made by using the FFT-NS-2 algorithm in MAFFT v7.407 program, and three regions (50~246aa; RBD domain; 600~796aa) were picked up. Finally, three phylogenetic trees were constructed using the Molecular Evolutionary Genetics Analysis (MEGA) software (version 10.0.5) by the maximum likelihood method with the bootstrap tests (100 replicates), and Fasttree software (version 1.43) were used to visualize these trees.

#### SNP recognition and comparison

The SNPs of each sequence were defined as the sites variant from the reference sequence. The ancestral sequence of the phylogenetic tree was used as the reference sequence, which was estimated by python package TimeTree using Jukes-Cantor model and default settings. Site with the unsolved nucleotide N and gap was ignored. The site number of the SNP was decided by sequentially combined the CDS sequences in the whole genome. The phylogenetic tree and the matching SNP for the tips was plotted by ggtree. The geo-distribution and genotype of strains with SNP sites in Wuhan and other areas were plotted by ggplot2. The SNP sites of 2019-nCoV were compared with SARS-like virus in one bat and eight pangolin genome sequences. The pangolin sequences EPI_ISL_410543 and EPI_ISL_410544 were removed for too many unsolved nucleotides. The adjacent nucleotide of the two SNP sites 8517 and 27641 were extracted from the alignment. The translation-reading frame was inferred by using the GenBank annotations for the EPI_ISL_402125 strain.

#### Tracking individual shared mutations

The divergent evolution of G1 and G2 was tracked with the help of shared mutations as following steps. First, we removed the substitution that appeared only once among all the genomes to avoid random mutations or sequencing errors that might mislead the inference. Second, genomes in G1 and G2 were grouped by the number of varied nucleotides compared with the reference sequence of G1 (EPI_ISL_402125) or G2 (EPI_ISL_406801). In total, we divided all the genomes in G1 and G2 separately into three groups with 1, 2 and 3 varied nucleotides. Third, the strains in groups with 1 varied nucleotides were plotted in the tree with branch length of 1, then, strains with 2 varied nucleotides were plotted in the tree with branch length of 2, it should be noted that the strains with substitution site contained the site in group 1 was plotted behind the branch with responding site. Similar ways were used to group 3.

#### Structure prediction and analysis

Based on the computer-guided homology modeling method, the structural models were constructed by SWISS-MODEL online server^33^. The model of nCoV 2019 Spike protein used the Cryo-EM structure of SARS coronavirus S-protein (PDB ID: 6ACD)^34^ as the template.

---

**Supplemental Figure 1.**
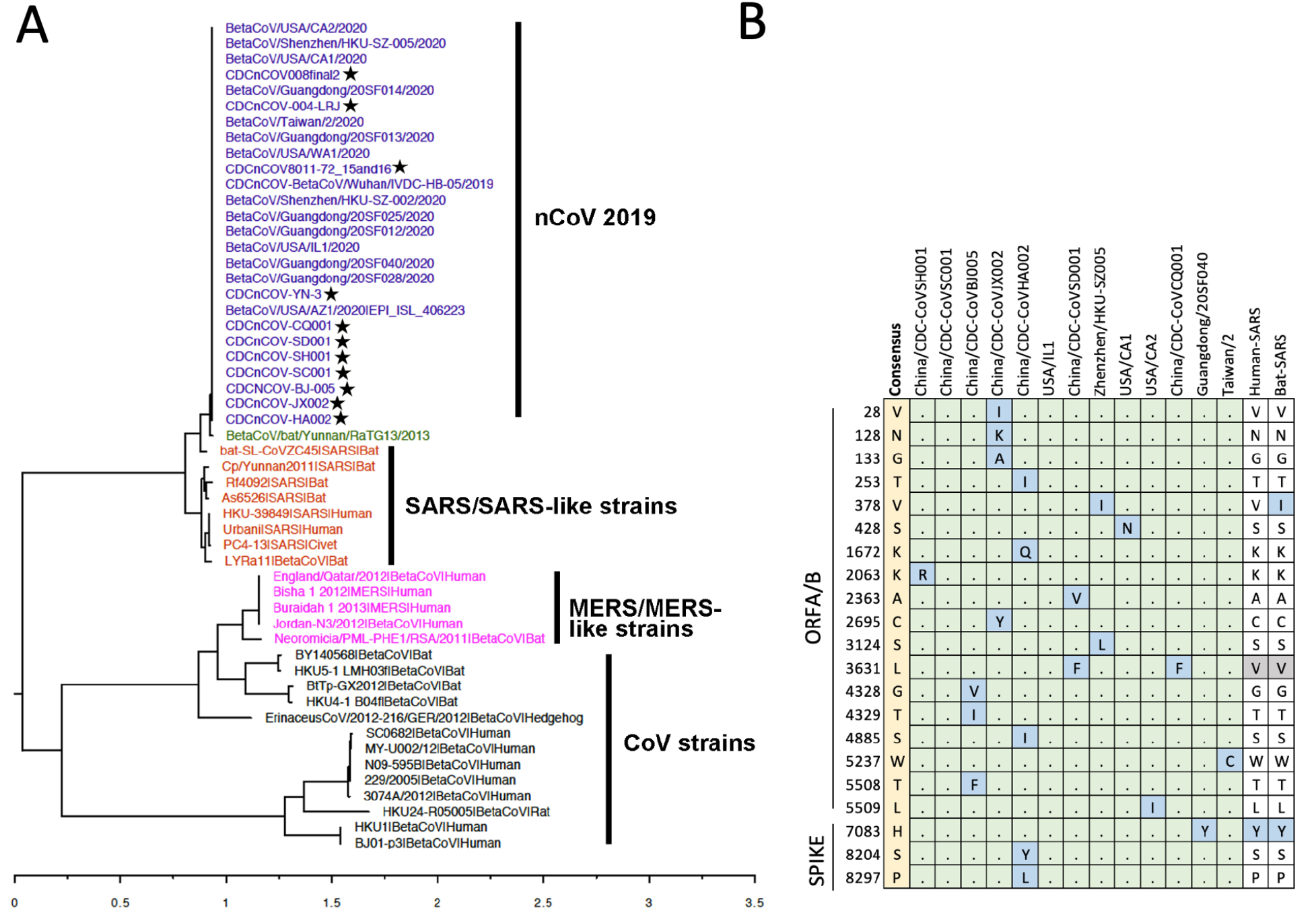
**(A)** Phylogenetic analysis of 46 full-length translated genomes of 2019-nCoV and 27 beta coronaviruses available in the public domain were aligned and estimated using MEGA7. 11 newly reported nCoV strains were highlighted using stars. **(B)** Amino acid substitutions within the same set of genomes were identified, and compared to consensus sequence of Human, and Bat-SARS.

**Supplemental Figure 2.**
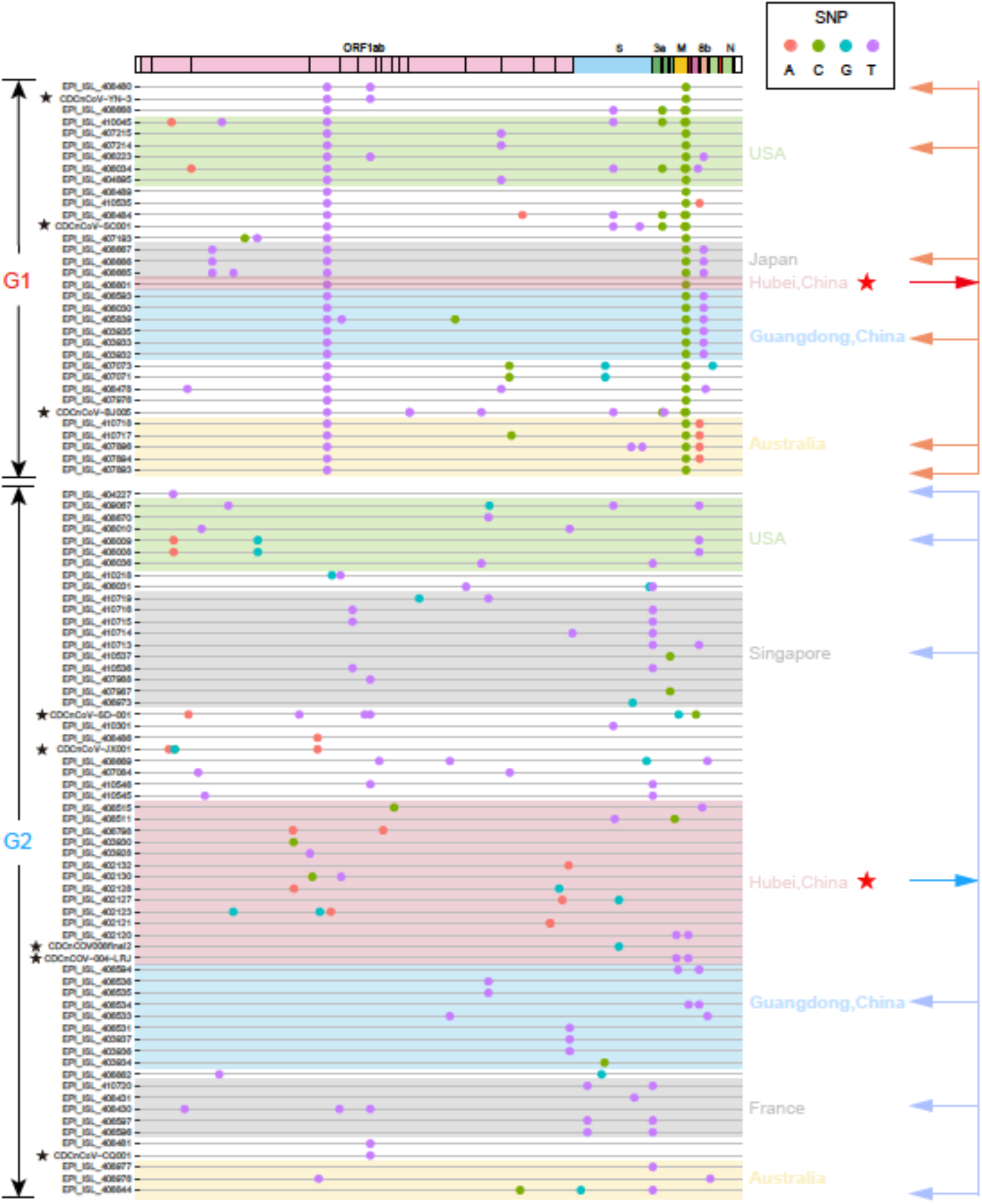
Co-circulation of two SARS-CoV-2 genotypes from Wuhan to other regions. The genomes of groups G1 and G2 are grouped individually by their collection locations. Five sampled locations are highlighted in different colours. SNPs are shown in small circles.

**Supplemental figure 3.**
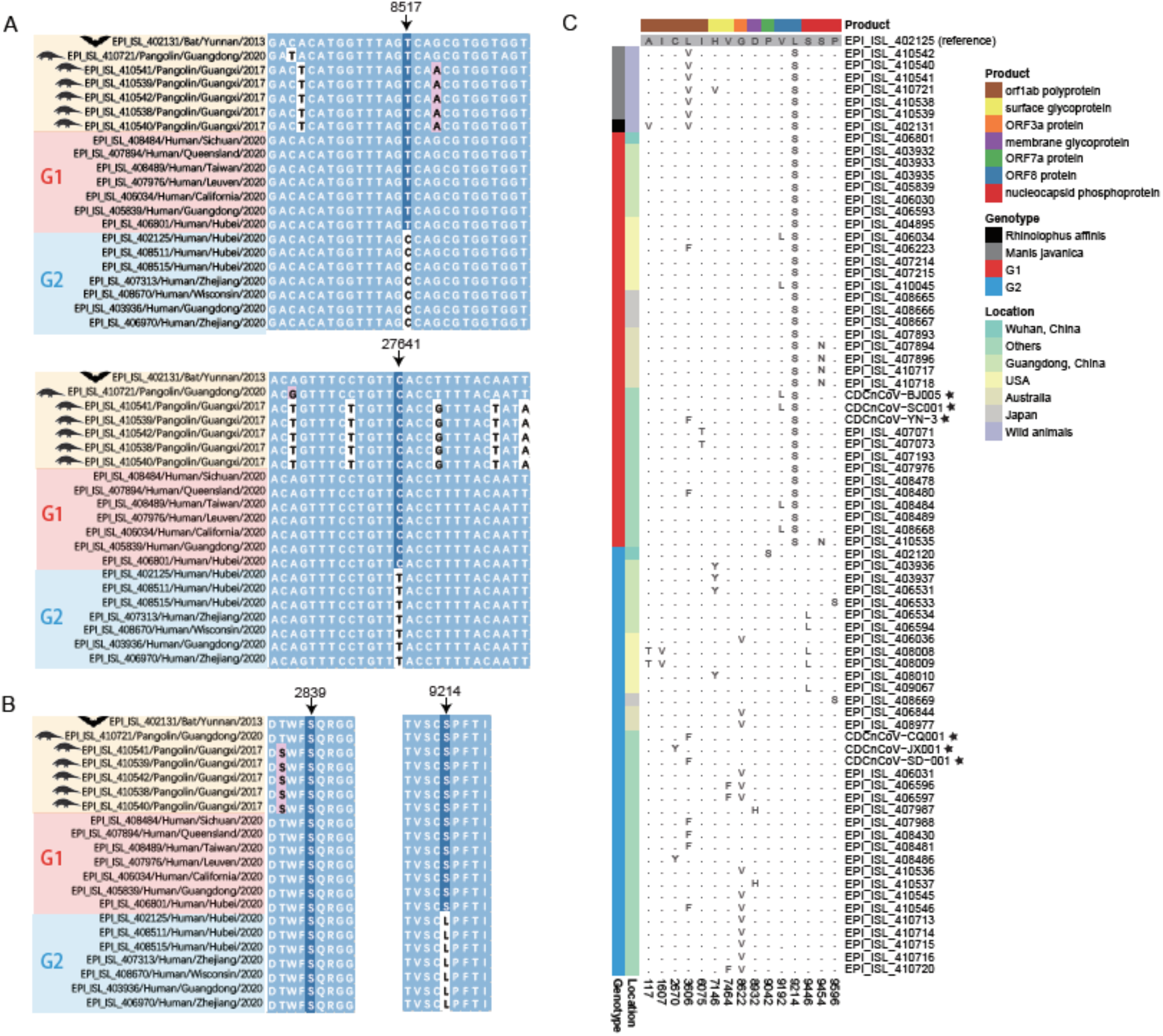
Major SNPs and amino acid substitutions among SARS-CoV-2 and similar coronaviruses isolated from bat and pangolin. **(A)** Two SNP markers in SARS-CoV-2 and similar coronaviruses isolated from bat and pangolin. **(B)** Comparison of two other minor SNPs. **(C)** Amino acid substitutions caused by non-synonymous SNPs (substitutions that appeared only once among all the sample genomes were removed). Strain EPI_ISL_402125 is used as reference. The numbering positions are listed at the bottom.Encoded proteins are highlighted as coloured bars at the top. Amino acids are shown as single letter or dot when it is the same as the reference strain.

**Supplemental figure 4.**
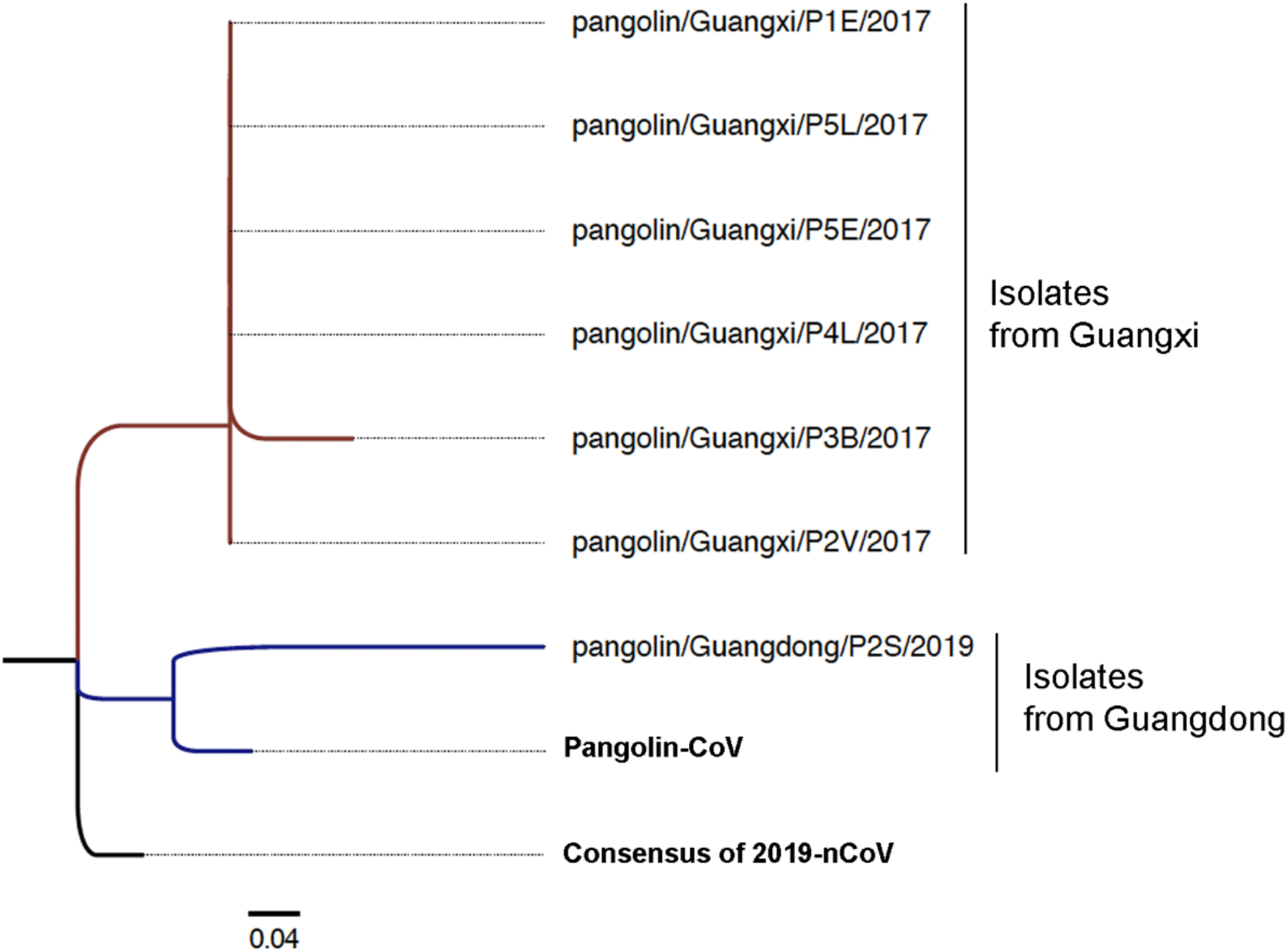
Phylogenetic analysis of the genome sequences from represented novel pangolin strains using the Maximum Likelihood method based on the JTT matrix-based model on MEGA v7.0.

**Supplemental Fig 5.**
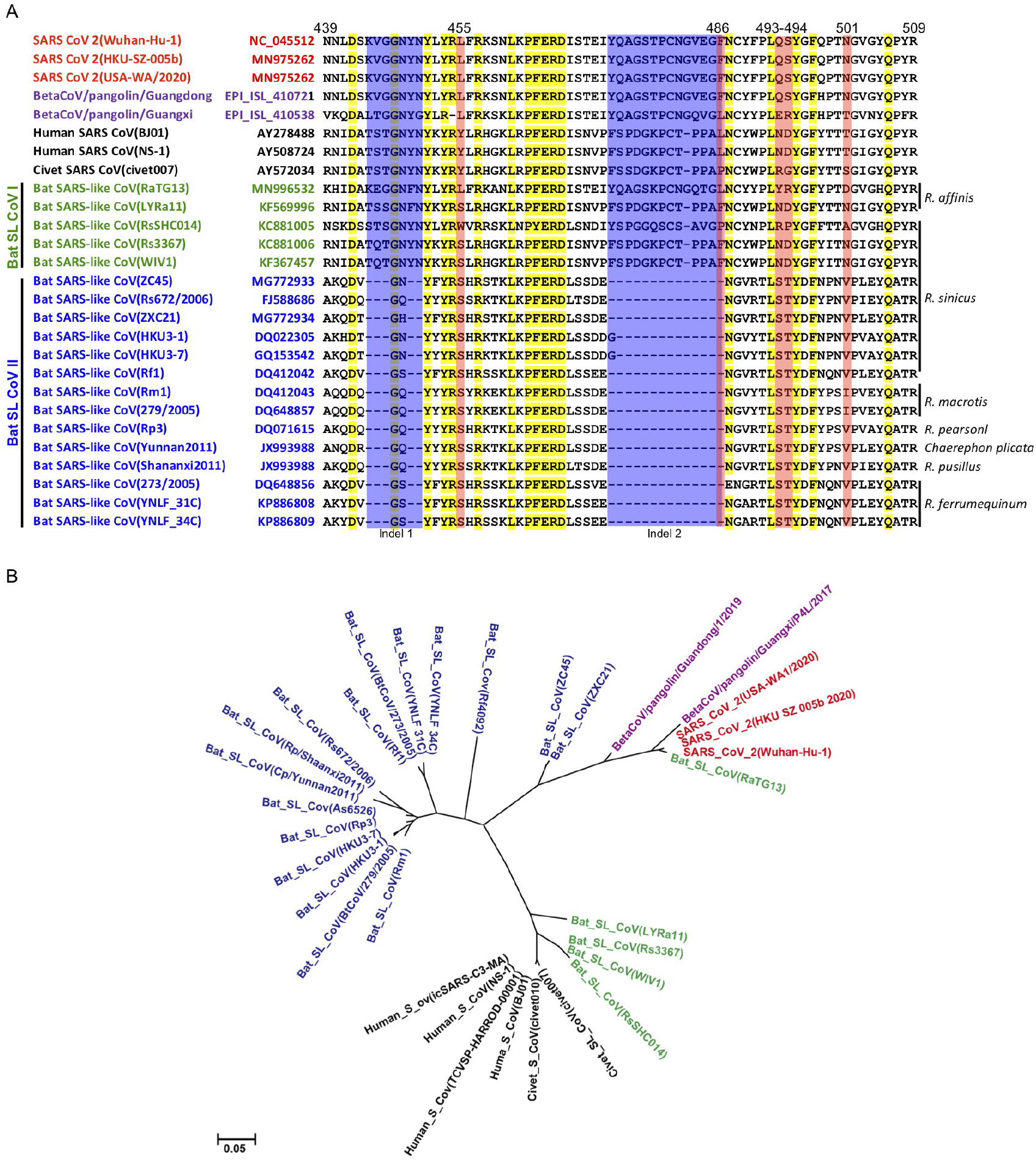
**(A)** Phylogenetic analyses of amino acid sequences of 2019-nCoV, human/civet SARS CoV and Bat SARS-like CoV from different bat species. The evolutionary history was inferred by using the Maximum Likelihood method based on the JTT matrix-based model conducting in MEGA7 software. The scale bar represents the number of substitutions per site. **(B)** Multiple alignment of the amino acid sequences of the receptor-binding motifs of the spike proteins of 2019-nCoV and SARS CoV and the corresponding sequences of bat SARS-like CoVs in different *Rhinolophus* species. Highly conserved residues are highlighted in yellow. Amino acid deletions in some bat SARS-like CoVs are labeled with blue. The five critical residues for receptor binding in 2019-nCoV at positions 455, 486, 493, 494 and 501(corresponding to 442,472,479,487,491 human/civet SARS CoV respectively) are highlighted in pink.

**Supplemental Fig 6.**
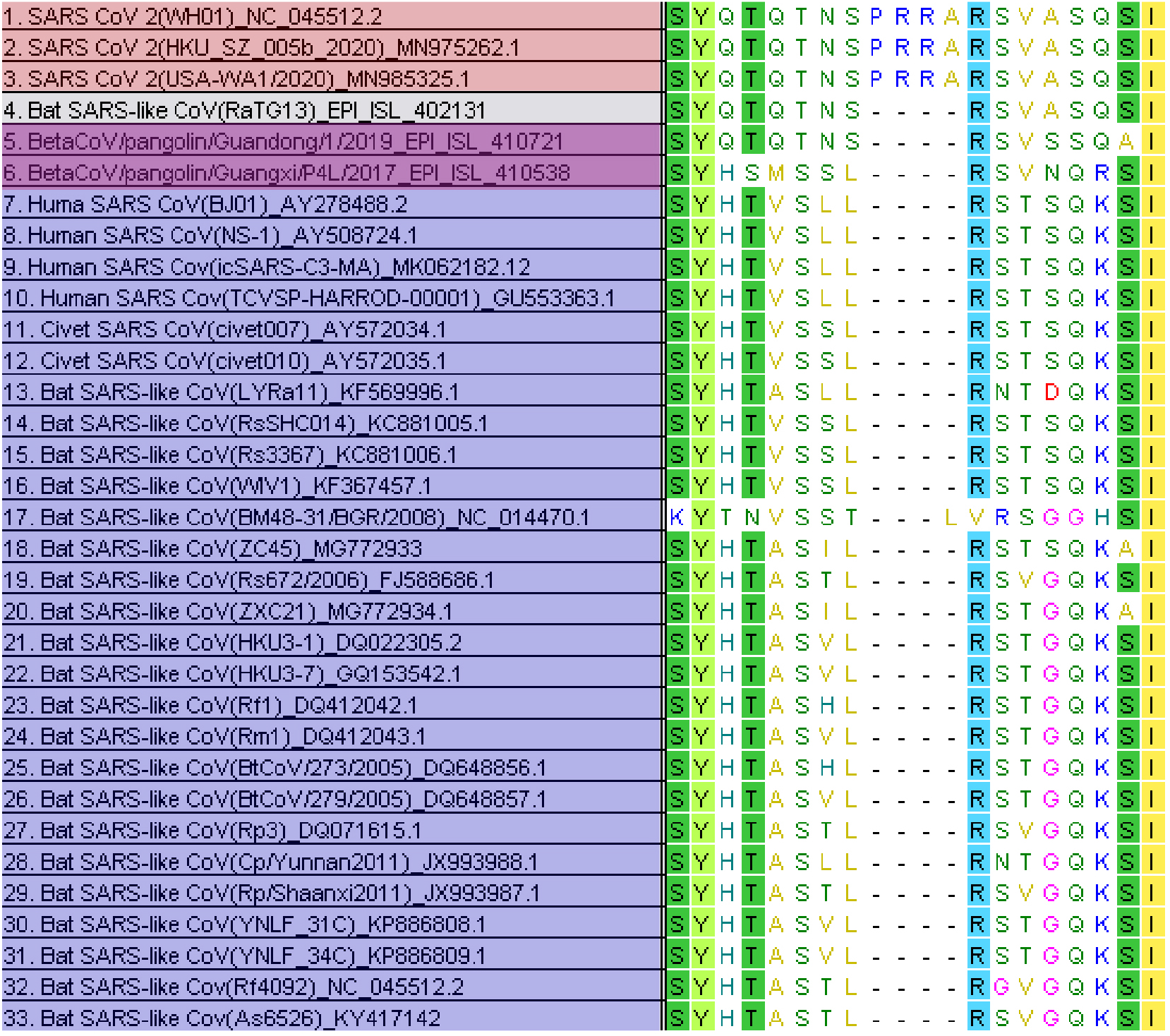
Unique insertion of a potential furin or TMPRSS2 cleavage site between S1 and S2 domain of the 2019-nCoV spike protein. (A) Insertion mutation at amino acid position 681 of 2019-nCoV spike protein and its sequence comparison with human and bat SARS-like CoVs. The multiple alignment was conducted by using the Clustal in MEGA7 software with default parameters.

**Supplemental Table 1:** Sequences of Bat SARS-like CoV Clade I collected from NCBI

| NO. | Name of Strains | Accession Number | Host species | Collection Date |
| --- | --- | --- | --- | --- |
| 1 | Bat SARS-like coronavirus isolate Rs9401 | KY417152 | *Rhinolophus sinicus* | 30-Dec-2016 |
| 2 | Bat SARS-like coronavirus isolate Rs7327 | KY417151 | *Rhinolophus sinicus* | 30-Dec-2016 |
| 3 | Rhinolophus affinis coronavirus isolate LYRa3 | KF569997 | *Rhinolophus sinicus* | 20-Aug-2013 |
| 4 | Bat SARS-like coronavirus Rs3367 | KC881006 | *Rhinolophus sinicus* | 08-Apr-2013 |
| 5 | Bat SARS-like coronavirus WIV1 | KF367457 | *Rhinolophus sinicus* | 08-Jul-2013 |
| 6 | Bat SARS-like coronavirus isolate Rs4874 | KY417150 | *Rhinolophus sinicus* | 30-Dec-2016 |
| 7 | SARS-like coronavirus WIV16 | KT444582 | *Rhinolophus sinicus* | 19-Aug-2015 |
| 8 | Bat SARS-like coronavirus isolate Rs4231 | KY417146 | *Rhinolophus sinicus* | 17-Apr-2013 |
| 9 | Bat SARS-like coronavirus RsSHC014 | KC881005 | *Rhinolophus sinicus* | 17-Apr-2011 |
| 10 | Bat SARS-like coronavirus isolate Rs4084 | KY417144 | *Rhinolophus sinicus* | 18-Sep-2012 |
| 11 | Rhinolophus affinis coronavirus isolate LYRa11 | KF569996 | *Rhinolophus affinis* | 22-Aug-2013 |
| 12 | Bat coronavirus isolate RaTG13 | MN996532 | *Rhinolophus affinis* | 24-Jul-2013 |

**Supplemental Table 2:**
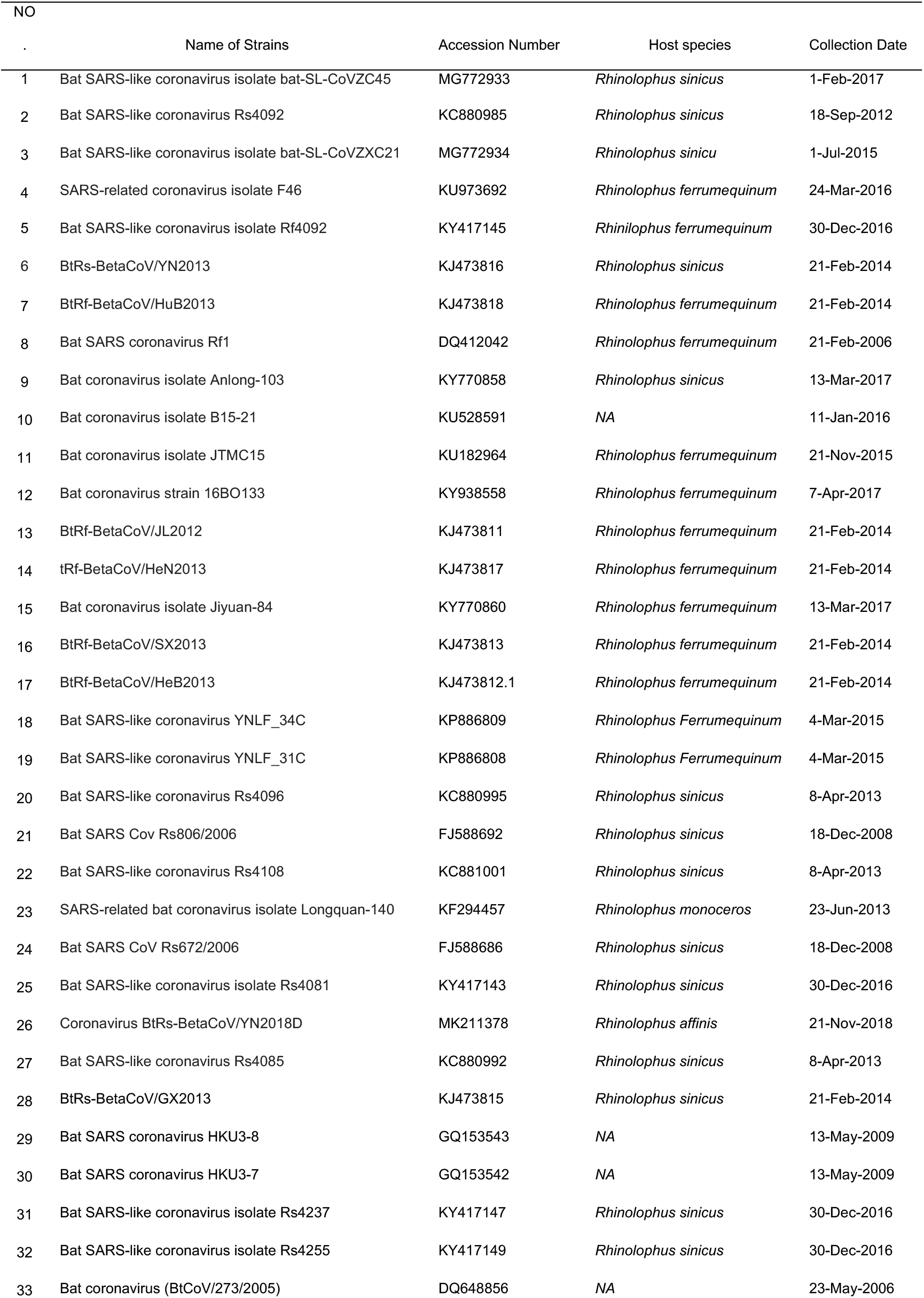
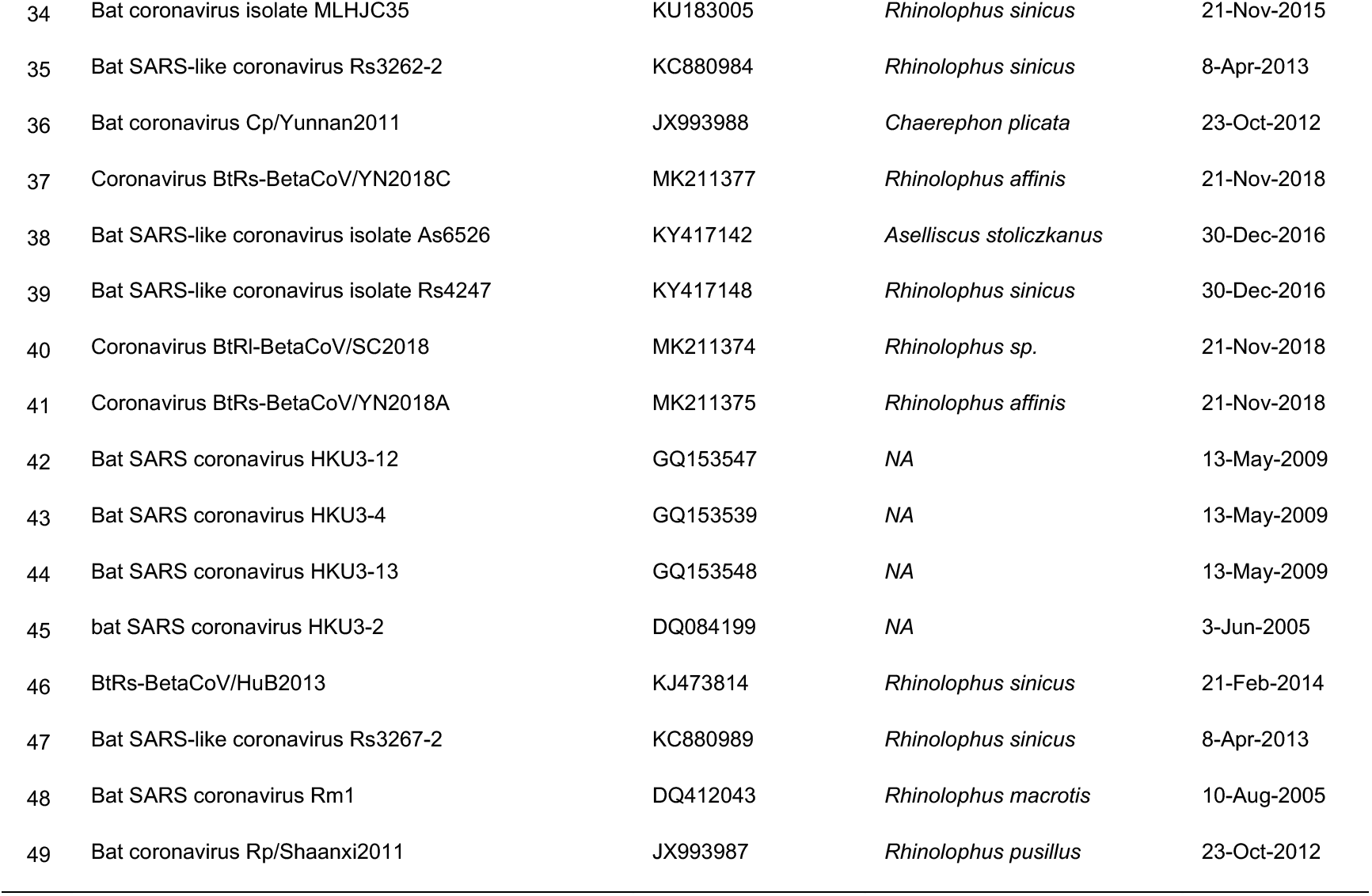
Sequences of Bat SARS-like CoV clade II collected from NCBI

